# Generalized locomotion of brittle stars with a flexible number of arms

**DOI:** 10.1101/616383

**Authors:** Daiki Wakita, Katsushi Kagaya, Hitoshi Aonuma

**Affiliations:** Graduate School of Life Science; Hokkaido University; Sapporo, Hokkaido, 060-0810; Japan; The Hakubi Center for Advanced Research; Kyoto University; Yoshida-Konoe, Kyoto, 606-8501; Japan; Seto Marine Biological Laboratory, Field Science, Education and Research Center; Kyoto University; Shirahama, Wakayama, 649-2211; Japan; Research Institute for Electronic Science; Hokkaido University; Sapporo, Hokkaido, 060-0812; Japan

**Keywords:** echinoderm, radial symmetry, limb number, moving direction, Bayesian statistical modeling, WAIC

## Abstract

Typical brittle stars have five radially symmetrical arms, which coordinate to move their body in a certain direction. However, some species of them show individual difference in the number of arms. We found this trait unique since intact legged animals each own a fixed number of limbs in general. How does a single species manage such different numbers of motile organs to realize adaptive locomotion? We here described four-, five-, six-, and seven-armed locomotion with the aim to generalize a common rule which is flexible with arm numbers in brittle stars. We mechanically stimulated an arm in *Ophiactis brachyaspis* to analyze escape direction and arm movements. Gathering quantitative indices and employing Bayesian statistical modeling, we figured out an average locomotion: regardless of the total number of arms, a front position emerges at one of the second neighboring arms to a mechanically stimulated arm, while side arms adjacent to the front synchronously work as left and right rowers. We suggest a model where some afferent signal runs either clockwise or anticlockwise along the nerve ring while linearly counting how many arms it passes. This idea would explain how ‘left and right’ emerges in a radially symmetrical body via a decentralized system.

## Introduction

Legged animals utilize appendages to move around on the ground. In most cases, intact adults of each species have a constant number of motile organs, such as four in many mammals and six in most insects. Supposedly, each species adopts a number-specific mechanism of locomotion. In this context, some species of brittle stars (Echinodermata: Ophiuroidea) exhibit an appealing individual difference, where some intact individuals have five appendages or less, while others have six or more (Fig. 1). This difference in number is usually found in fissiparous species, which undergo asexual reproduction by fission and regeneration (Boffi, 1972; Mladenov et al., 1983; Mladenov and Emson, 1984).

**Fig. 1.**
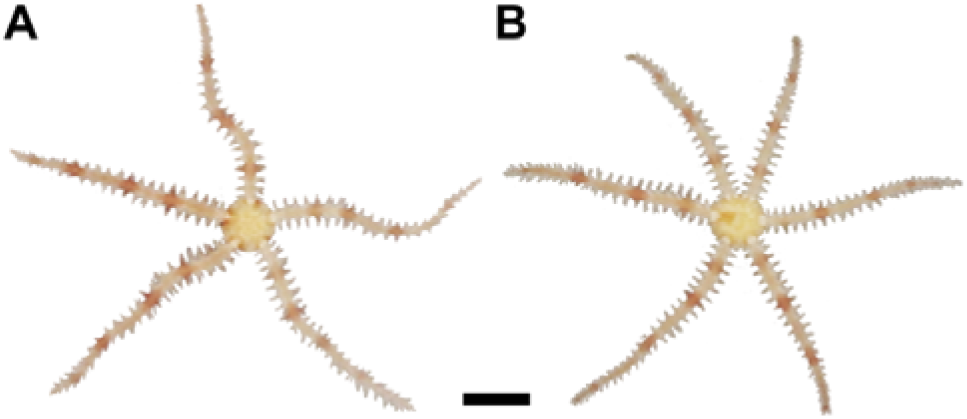
The fissiparous brittle star *Ophiactis brachyaspis*. A: a five-armed individual. B: a six-armed individual. The scale bar represents 2 mm.

As typical echinoderms show pentaradial symmetry, most ophiuroid species standardly have five multi-jointed appendages called “arms,” which extend from the “disk” at the center. Previous studies have described arm movements in the locomotion of five-armed species in qualitative terms (Romanes and Ewart, 1881; Preyer, 1887; von Uexküll, 1905; Glaser, 1907; Arshavskii et al., 1976a,b; Clark et al., 2019) as well as in quantitative terms (Astley, 2012; Kano et al., 2017). Several locomotor modes have been known even in a single species. An often reported mode, referred to as “breast stroke” (Arshavskii et al., 1976a,b) or “rowing” (Astley, 2012), is characterized by a leading arm facing forward, two side arms working as left and right rowers, and two back arms dragged passively (Romanes and Ewart, 1881; Preyer, 1887; Glaser, 1907; Arshavskii et al., 1976a,b; Astley, 2012; Kano et al., 2017). Some studies have observed another locomotor mode, called as “paddling” (Arshavskii et al., 1976a) or “reverse rowing” (Astley, 2012), where a backmost arm is dragged while the other four actively row (Preyer, 1887; von Uexküll, 1905; Glaser, 1907; Arshavskii et al., 1976a; Astley, 2012). The ophiuroid body creeps in a certain direction with such bilaterally coordinated manners (Astley, 2012). Since the ‘role’ of each arm switches when the body changes moving direction (Arshavskii et al., 1976a; Astley, 2012), brittle stars do not have consistent antero-posterior and left-right axes in behavior.

In neurological aspects, the ophiuroid nervous system principally comprises a circumoral nerve ring in the disk and radial nerve cords running into each arm (Cobb and Stubbs, 1981, 1982; Ghyoot et al., 1994; Bremaeker et al., 1997; Zueva et al., 2018). At each branch to the radial nerve, the nerve ring includes regional concentrations of neural cell bodies, which are recognizable as ganglia controlling some organs (Ghyoot et al., 1994). Some behavioral studies have supported the essential roles of the nerve ring in locomotion. Menthol-anesthetic experiments indicated its function in initiating locomotion (Matsuzaka et al., 2017). Nerve cut experiments have demonstrated its necessity for coordinating arms (Mangold, 1909; Diebschlag, 1938; Arshavskii et al., 1976a,b; Clark et al., 2019).

Although the five-armed locomotion in common brittle stars and the individual difference in specific species have been viewed in different contexts, none has combined them to spotlight ophiuroid locomotion across different numbers of symmetrical repeats. Some studies have observed locomotion with several arms cut (Arshavskii et al., 1976b; Kano et al., 2017; Matsuzaka et al., 2017; Clark et al., 2019), but pentaradial architecture remains at the disk and thus at the nerve ring. On the other hand, when structural divisions at the center differ in count, the nerve ring must have a different number of branches to radial nerves—implying another number of ganglia. Given the reported functional importance of the nerve ring in locomotion, such plus or minus in the central network seems to bring a huge issue in integrating an individual.

How does an animal species realize adaptive locomotion while managing a changeable number of motile organs and network branches? The aim of our study is to generalize ophiuroid locomotion which is flexible with the number of radially symmetrical arms. We targeted four-, five-six-, and seven-armed intact individuals in a fissiparous species, *Ophiactis brachyaspis* Clark, 1911 (Fig. 1). For probing into inter-arm communication, we gave an arm an aversive tactile stimulus to observe the reactions of the other arms. Although electrophysiological approaches are difficult in the small body of fissiparous ophiuoids, we can suppose each arm’s movement is a simple reflection of neural activity, so external behavioral modeling will infer internal neural networks. As a primary hypothesis, brittle stars would have a decentralized neural network, where each functional unit affects nearest-neighbor units while ignoring distant ones to allow the flexibility in total number. With such a design, one possible expectation is that a threat to an arm makes its both neighbors push toward the stimulus, so that the disk can escape in the opposite direction to the stimulus.

In this study, we quantitatively described post-stimulus locomotion based on arm movements and escape direction, employing Bayesian estimation and model selection to understand their potential structures as reasonable distributions. Our results indeed reinforced the support of a decentralized strategy in the ophiuroid body; however, escape direction did not always frequent the opposite of stimulation, contrary to our initial expectation. We suggest a generalized model as follows: a threat to an arm makes an afferent signal which asymmetrically dominates either of clockwise or anticlockwise detour of the nerve ring; in a decided detour, the stimulated arm’s *second* neighboring arm is highly probable to be a leading arm, with the leader’s side arms working as left and right synchronous rowers. Thus regardless of the total number of arms, ophiuroid locomotion shows a common anterior pattern, which could be positioned by linearly counting how many arms some signal passes in one direction along a circular pathway. We also provide a unique idea of how ‘left and right’ emerges in a multidirectional body to make a unidirectional behavior.

## Materials and methods

### Animals

We used the fissiparous brittle star *Ophiactis brachyaspis* (Fig. 1). In nature, this species densely inhabits the upper and side surfaces of rough rocks or other adherent organisms such as sponges. Some of its arms lie in interstices while some rise from the substrate; suspension feeding ophiuroids show such a posture to capture particles (Warner, 1971). Animals collected in Shirahama Aquarium, Kyoto University, were reared in a laboratory aquarium (600 × 600 × 600 mm) filled with artificial seawater at 25–28°C with the salinity of 32–35‰ (TetraMarin Salt Pro, Tetra Japan Co, Tokyo, Japan). Body size ranged 1.5–3.0 mm in disk diameter and 5–15 mm in arm length. The number of arms was six in the majority—about 70%—and five for the others. Four- and seven-armed bodies each were found only in one individual.

### Behavioral experiments

To investigate locomotion, we chose 10 five-armed individuals and 10 six-armed individuals, in each of which the lengths of arms did not differ by more than twice (c.f. Fig. 1). Four- and seven-armed individuals were also targeted; we obtained one for each. Each individual was put in a horizontal flat acrylic case (105 × 75 × 22 mm) filled with 100 mL of artificial seawater from the laboratory aquarium. There were no strong light gradient and no strong wind. Locomotion was recorded in aboral view using a digital camera (EOS8000D, Canon, Tokyo, Japan) with videos saved in MP4 format. We applied tactile stimuli to arm tips using a toothpick. Stimulating an arm with subsequent observation was defined as one trial. The next trial came at the anticlockwise neighboring arm with an interval of more than two minutes. With repeating this rotation in order, every arm was stimulated at least three times for each individual.

### Measurements

Per five- or six-armed individual, we analyzed three trials which showed the longest moving distances of the disk. In the four- and seven-armed cases, we picked out 15 trials with the longest moving distances. Analyzed duration for each trial was one minute after beginning to move the disk following each stimulus (c.f. Videos S1, S2). The stimulated arm in each trial was numbered 1, which was followed anticlockwise by the other arms; *α* is the index of arms (*α* = 1, 2, 3, 4, 5 in the five-armed instance). We tracked two coordinate points of the *α*-th arm using a tracking software Kinovea ver. 0.8.27 (http://www.kinovea.org/, accessed 4 December 2018) at 10 f.p.s.: *P*_*α*_(*t*) = (*x*_*α*_(*t*), *y*_*α*_(*t*))—the attachment point between the α-th arm and the disk viewed aborally—and *P*′_*α*_(*t*) = (*x*′_*α*_(*t*), *y*′_*α*_(*t*))—the point at half the length of the *α*-th arm, in terms of the range from the center of disk to the arm tip—at the *t*-th frame (Fig. S1). For the latter, we did not choose each arm’s tip because it often rose and showed casual movements seeming irrelevant to locomotion as Matsuzaka et al. (2017) indicated. *P*_cent_(*t*) was defined as the center of gravity of all arms’ *P*_*α*_(*t*) (Fig. S1). The *α*-th arm’s length (*L*_*α*_) was defined as the maximum length of the segment *P*_*α*_(*t*)*P*′_*α*_(*t*) in the analyzed duration. Note that *L*_*α*_ is a variable sampled in each trial, not accounting for the constant length of each arm. Moving distance (*S*) was measured as the length of *P*_cent_(1)*P*_cent_(*T*), where *T* is the total number of frames, i.e. 600 (Fig. S1). We assessed moving direction (*Θ*) as follows:

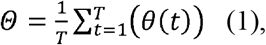

where *θ*(*t*) is the angle made by the two segments *P*_c_(1)*P*_c_(*T*) and *P*_1_(*t*)*P*_cent_(*t*) (Figs S1, S2). *Θ*, which takes the range from −180 to 180 deg, is 0 deg when the disk moves in the opposite direction to the stimulated arm. A negative or positive value of *Θ* represents that the disk movement is angled clockwise or anticlockwise, respectively, from the opposite direction to the stimulated arm. For later uses in statistics, the dummy variable *Θ*_sign_ is defined as

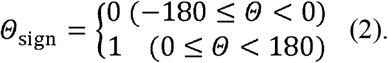

The segment *P*_cent_(*t*)*P*′_*α*_(*t*) during locomotion swung around *P*_cent_(*t*)*P*_*α*_(*t*), so the *α*-th arm’s angle at the *t*-th frame (*φ*_*α*_(*t*)) was defined as the angle made by these two segments (Fig. S1). *φ*_*α*_(*t*) is negative or positive when *P*_cent_(*t*)*P*′_*α*_(*t*) is angled clockwise or anticlockwise, respectively, from *P*_cent_(*t*)*P*_*α*_(*t*). *φ*_*α*_(*t*)’s angular velocity (*ω*_*α*_(*t*)) was calculated with a five-point moving average method, and then smoothened with a low-pass filter with the cutoff frequency of 1.0 Hz (Fig. S2). To quantify to what extent each arm functions as left or right rower, we focused on that returning was faster than pushing in rowing arms. The filtered *ω*_*α*_(*t*) was thus analyzed to evaluate the degree of a leftward or rightward bias in movement, which is represented by *B*_*α*_(named after “bias”; Fig. S2):

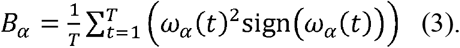

Assuming that a directional bias results from a speed difference between pushing and returning in each arm, we can rephrase *B*_*α*_ as the α-th arm’s degree of being a left or right rower. A largely negative value of *B*_*α*_ represents that the *α*-th arm moves clockwise faster than anticlockwise, indicating that it slowly pushes leftward and fast returns rightward viewed proximally from the disk. On the contrary, *B*_*α*_ is largely positive when the *α*-th arm pushes rightward (clockwise). Its value is close to zero when the *α*-th arm pushes leftward and rightward equally or is dragged without actively returning. We also extracted frequency components in the non-filtered *ω*_*α*_(*t*) of each arm using Fourier transforms. *F*_*α*_ was defined as the frequency at the peak amplitude in the *α*-th arm.

Besides for *B*_*α*_, we used the filtered *ω*_*α*_(*t*) to calculate Kano et al.’s (2017) “*E*_*ij*_,” namely, the degree of synchronization between two arms:

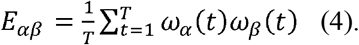

A negative or positive value of *E*_*αβ*_ represents that the movements of the *α*- and *β*-th arms synchronize in the opposite or same direction, respectively. A value around zero represents that the two arms move without strong correlation or are static.

### Statistical modeling

We built statistical models for later comparative assessments with the following procedure. Firstly, to examine the structure of a possible bimodality in moving direction, we assume that *Θ* is subjected to a single von Mises distribution (*f*_vM_, ‘circular normal distribution’),

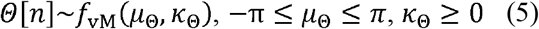

or a mixture of two von Mises distributions,

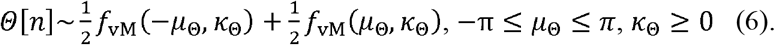

Hereafter, *n* takes one to the total number of trials, so that *Θ*[*n*] denotes the *n*-th element of *Θ*. The parameters as random variables *μ*_*Θ*_—converted to radians for modeling—and *κ*_*Θ*_ are analogous to the mean and the reciprocal of variance, respectively, in normal distribution. For the mixed case, we assume that the two distributions are symmetrical to each other with respect to the position of 0 deg.

Secondly, to understand what brings a trial-by-trial variability of *B*_*α*_, we parametrize *L*_*α*_, *S*, *Θ*, *Θ*_sign_, and *F*_*α*_ each as an explanatory variable for *B*_*α*_. We assume the normal distribution *f*_norm_(*μ*, *σ*), where *μ* and *σ* respectively represent the mean and standard deviation (s.d.), as follows:

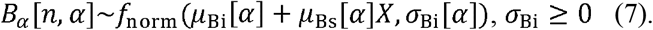

Here, *μ*_Bi_, *μ*_Bs_, and *σ*_Bi_ are arm-by-arm parameters and *X* is an explanatory variable to which *L*_*α*_[*n*, *α*], *S*[*n*], *Θ*[*n*], *Θ*_sign_[*n*], or *F*_*α*_[*n*, *α*] is assigned. *S*, *Θ*, and *Θ*_sign_ are common values for all the arms in the same trial. The categorical index *Θ*_sign_ is to know whether *B*_*α*_ varies continuously by *Θ* or switches discretely by the sign of *Θ*. In this instance, *μ*_Bs_ represents the mean’s difference between the negative and positive cases since this variable disappears when *Θ*_sign_ is zero (−180 ≤ *Θ* < 0) and appears when *Θ*_sign_ is one (0 ≤ *Θ* < 180). The model without the member *μ*_Bs_[*α*]*X*, i.e. without an explanatory variable, is for comparison. In parallel, let us consider whether *B*_*α*_ is better explained by individuality, namely, the quality made by some individual difference other than arm number as to five- and six-armed animals. Consideration of individuality is given to the mean’s intercept *μ*_Bi_:

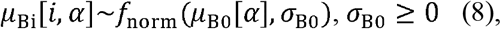

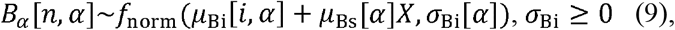

where *i* takes one to the total number of individuals (i.e. 10) and the hyperparameters *μ*_B0_ and *σ*_B0_ are random variables. Let *σ*_B0_, which is common in all arms, have a weakly informative prior as

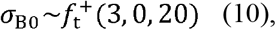

where 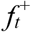 denotes the half *t* distribution and the parenthetical parameters represent the degree of freedom (*v*), location (mean when *v* > 1), and scale (s.d. divided by 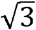 when *v* = 3), respectively.

The final modeling is to examine which of *Θ* and *Θ*_sign_ is a better explanatory variable for *E*_*αβ*_ in four- to seven-armed animals:

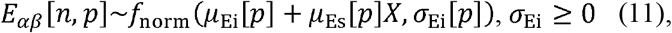

where *μ*_Ei_, *μ*_Es_, and *σ*_Ei_ are pair-by-pair parameters and the explanatory variable *X* takes *S*[*n*], *Θ*[*n*] or *Θ*_sign_[*n*]. Also considered is the model without the explanatory member *μ*_Es_[*p*]*X*.

Employing the Bayesian approach, posterior distribution of each parameter was estimated by the no-U-turn sampler (NUTS) (Hoffman and Gelman, 2014)—a variant of Hamiltonian Monte Carlo (HMC) algorithm. In each sampling, we totally obtained 10,000 NUTS samples from four Markov chains, in each of which every 40th generation was sampled in 100,000 iterations after a warmup of 5,000, with the target acceptance rate of 0.8. Convergence of each parameter was checked by trace plots, the potential scale reduction factor 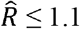, and the effective sample size 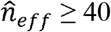, i.e. at least 10 per chain (Gelman et al., 2013). We assessed the predictability of the models based on WAIC (Watanabe, 2019, 2010), as this criterion is applicable to our models containing mixed distributions (Eqn 6) or hierarchical parameters (Eqns 8, 9). We developed the resultant statements according to better predicting models, which yielded smaller WAICs than the others considered. For comparison between the models, we referred to the difference as

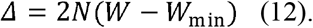

*N* is the total number of measured samples; multiplication by 2*N* is for the AIC scaling (Gelman et al., 2013). *W* is a given model’s WAIC while *W*_min_ is the smallest WAIC among those of the proposed models, so that Δ is zero in the best performed models. In presenting figures, the posterior predictive distributions of *Θ* are shown based on the parameters’ posterior distributions in a better performed model, each indicating a probability distribution which is expected to generate a random variable in a new trial. To visualize *B*_*α*_ and *E*_*αβ*_ dependent on a better explanatory variable, we obtained the median of each posterior distribution under a model including the explanatory variable not only in the mean but also in the s.d.; Eqn 7 or 9 was modified to

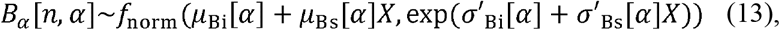

while Eqn 11 was replaced by

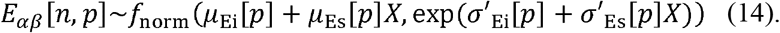

Exponentiation in scale is to make the s.d. positive while *σ*′_Bi_, *σ*′_Bs_, *σ*′_Ei_, and *σ*′_Es_ are random variables without constraints. We did not consider scale’s explanatory variables in WAIC comparing terms because the Markov chain simulation failed to converge in many cases. All statistical computation was performed in the software environment R ver. 3.5.1 (R Core Team, 2018), where Stan codes were compiled and executed by the R package “rstan” (Stan Development Team, 2018). All source codes and data are available from the Figshare repository (Wakita et al., 2019).

## Results

### Moving direction (*Θ*)

The measured data of the post-stimulus moving direction *Θ* (Figs S1, S2; Eqn 1 in “Materials and methods”) are shown in Fig. 2 by dot plots. For all the four-, five-, six-, and seven-armed cases, the model assuming a mixture of two von Mises distributions in *Θ* yielded smaller WAICs—better predictabilities—than that assuming a single distribution (Table 1). Compared with the small Δ (difference from the best one; Eqn 12 in “Materials and methods”) of the one-distribution model in four- and five-armed animals, the six- and seven-armed Δ values were larger enough to interpret that bimodality was more obvious in more arms. Following the better model in terms of WAIC, we hereafter show the results on the assumption of two distributions for all the cases.

**Table 1.** WAICs of statistical models for *Θ*, *B*_*α*_, and *E*_*αβ*_.

| Model | Specification |  | Four-armed |  |  | Five-armed |  |  | Six-armed |  |  | Seven-armed |  |  |
| --- | --- | --- | --- | --- | --- | --- | --- | --- | --- | --- | --- | --- | --- | --- |
| | | | Rank | WAIC | $\Delta$ | Rank | WAIC | $\Delta$ | Rank | WAIC | $\Delta$ | Rank | WAIC | $\Delta$ |
| $\Theta$ | Distribution number | | | | | | | | | | | | | |
| 1 | one |  | 2 | 0.876 | 0.917 | 2 | 1.192 | 0.764 | 2 | 1.518 | 5.55 | 2 | 1.860 | 10.7 |
| 2 | two |  | 1 | 0.845 | **0\*** | 1 | 1.179 | **0\*** | 1 | 1.425 | **0\*** | 1 | 1.502 | **0\*** |
| $B_\alpha$ | Explanatory variable<sup>†</sup> | Individ-<br>uality<sup>†</sup> | | | | | | | | | | | | |
| 1 | no | no | 3 | 4.217 | 4.94 | 8 | 4.671 | 60.4 | 5 | 5.338 | 58.9 | 4 | 4.940 | 25.7 |
| 2 | no | yes | — | — | — | 9 | 4.676 | 61.9 | 10 | 5.352 | 64.0 | — | — | — |
| 3 | $L_\alpha$ | no | 1 | 4.176 | **0\*** | 10 | 4.679 | 62.9 | 9 | 5.350 | 63.2 | 5 | 4.947 | 27.1 |
| 4 | $L_\alpha$ | yes | — | — | — | 12 | 4.688 | 65.4 | 12 | 5.382 | 74.9 | — | — | — |
| 5 | $S$ | no | 5 | 4.267 | 10.9 | 7 | 4.670 | 60.3 | 7 | 5.338 | 59.2 | 3 | 4.938 | 25.2 |
| 6 | $S$ | yes | — | — | — | 11 | 4.680 | 63.1 | 11 | 5.360 | 67.0 | — | — | — |
| 7 | $\Theta$ | no | 2 | 4.187 | 1.29 | 1 | 4.469 | **0\*** | 2 | 5.191 | 6.25 | 2 | 4.827 | 2.03 |
| 8 | $\Theta$ | yes | — | — | — | 2 | 4.477 | 2.16 | 4 | 5.208 | 12.2 | — | — | — |
| 9 | $\Theta_{\text{sign}}$ | no | 4 | 4.230 | 6.41 | 3 | 4.501 | 9.43 | 1 | 5.174 | **0\*** | 1 | 4.818 | **0\*** |
| 10 | $\Theta_{\text{sign}}$ | yes | — | — | — | 4 | 4.505 | 10.8 | 3 | 5.193 | 6.89 | — | — | — |
| 11 | $F_\alpha$ | no | 6 | 4.271 | 11.3 | 5 | 4.640 | 51.0 | 6 | 5.338 | 59.1 | 6 | 4.955 | 28.8 |
| 12 | $F_\alpha$ | yes | — | — | — | 6 | 4.644 | 52.3 | 8 | 5.347 | 62.4 | — | — | — |
| $E_{\alpha\beta}$ | Explanatory variable<sup>†</sup> | | | | | | | | | | | | | |
| 1 | no |  | 1 | 3.951 | **0\*** | 4 | 4.327 | 25.6 | 4 | 4.440 | 42.0 | 3 | 4.294 | 9.30 |
| 2 | $S$ | | 4 | 3.974 | 4.14 | 3 | 4.321 | 22.0 | 2 | 4.413 | 17.5 | 4 | 4.320 | 25.7 |
| 3 | $\Theta$ | | 2 | 3.953 | 0.365 | 2 | 4.292 | 4.90 | 3 | 4.416 | 20.0 | 2 | 4.282 | 1.58 |
| 4 | $\Theta_{\text{sign}}$ | | 3 | 3.959 | 1.45 | 1 | 4.284 | **0\*** | 1 | 4.394 | **0\*** | 1 | 4.279 | **0\*** |

\* $\Delta = 0$ , bolded, indicating a best supportive model. <sup>†</sup>Not considered if “no”; otherwise, considered in the mean of normal distribution.

**Fig. 2.**
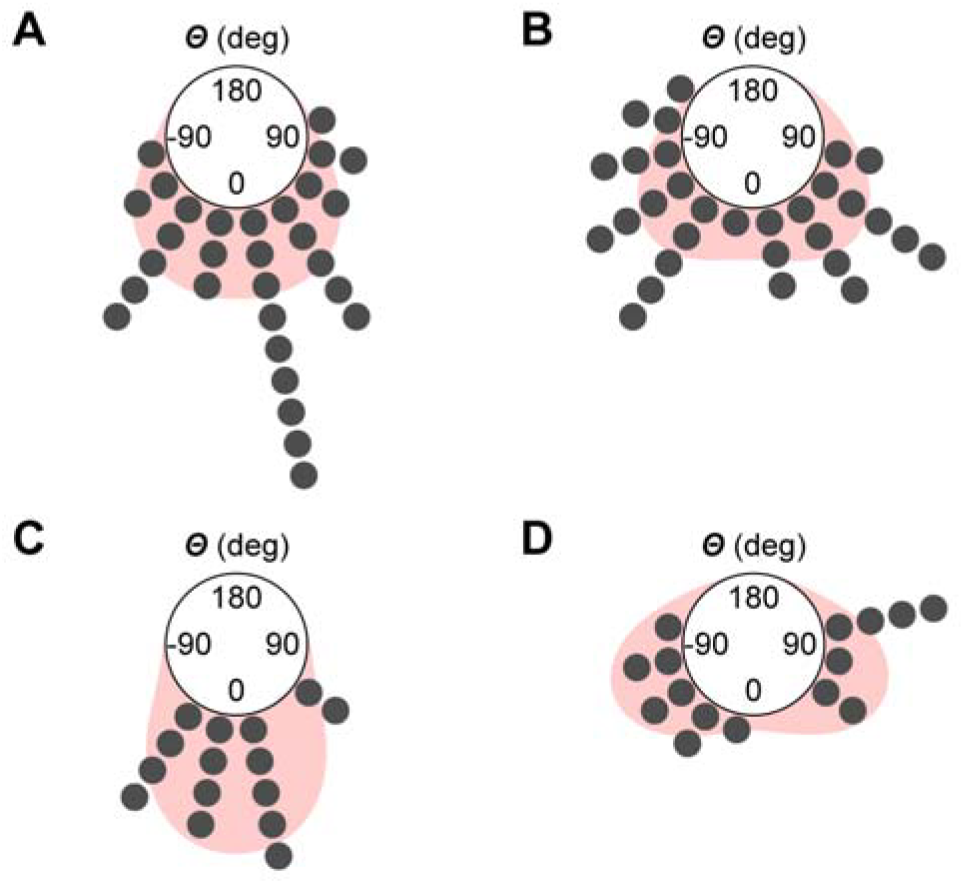
Circular plots of moving direction after aversive tactile stimulation in the brittle star *Ophiactis brachyaspis*. A: five-armed case (10 individuals, 30 trials). B: six-armed case (10 individuals, 30 trials). C: four-armed case (one individual, 15 trials). D: seven-armed case (one individual, 15 trials). The moving direction *Θ* is the measured angle based on the position of a mechanically stimulated arm (c.f. Figs S1, S2). *Θ* is 0 deg when the disk moves in the opposite direction to the stimulated arm, and is negative/positive when the disk movement is angled clockwise/anticlockwise, respectively, from the 0 deg. Each point represents *Θ* in each trial, which is grouped in a bin divided per 22.5 deg. Density plots on the background represent predictive distributions on the assumption of two symmetrical von Mises distributions.

The posterior medians of two distributions’ means, which were calculated separately for the negative and positive ranges, were ±17, ±29, ±46, and ±70 in four- to seven-armed animals, respectively. These estimated values signify that the more arms a brittle star had, the further two distributions of *Θ* were apart from each other (Fig. 2). In other words, the average moving direction of individuals with more arms was more angled from the opposite direction to the stimulated arm. Predictive distribution of *Θ* indeed depicted this trend (Fig. 2).

### Left or right rower (*B*_*α*_)

The measured data of *B*_*α*_—the *α*-th arm’s degree of being a left or right rower (Figs S1, S2; Eqn 3 in “Materials and methods”)—are schematized trial-by-trial in Figs S3–S6. As for the five- and six-armed populations, no-individuality models consistently took smaller WAICs than their counterparts where individuality was assigned to the mean of *B*_*α*_ (Table 1). We thus avoid mentioning individual difference within the same arm number.

Among *L*_*α*_, *S*, *Θ*, *Θ*_sign_, and *F*_*α*_, the five-armed *B*_*α*_ was better explained by the continuous moving direction *Θ*, whereas the six- and seven-armed cases rather emphasized its sign *Θ*_sign_ (Eqn 2 in “Materials and methods”) in discrete terms (Table 1). In four arms, the arm length *L*_*α*_ was chosen for a best explanatory variable although *Θ* showed a close performance. Given the dominance of the moving direction indicators as well as *Θ*’s bimodality (Fig. 2), we present the data of *B*_*α*_ separately by *Θ*_sign_—in which side moving direction angled from the midline of the stimulated arm. Two groups were here defined by whether it angled clockwise (*Θ*_sign_ = 0) or anticlockwise (*Θ*_sign_ = 1).

The *Θ*_sign_-based grouping exhibited a common locomotor mode among four-, five-, six-, and seven-armed animals in regards to *B*_*α*_’s posterior means. The directional property of each arm could be explained by how many arms we count from the stimulated arm. Primarily, one of the *first* neighboring arms to the stimulated arm consistently took the largest or second largest |*B*_*α*_|—absolute values of posterior means (Figs 3–6A,C). This *first* arm corresponds to the anticlockwise neighbor of the arm 1 when *Θ*_sign_ = 0 (Figs 3–6A) and the clockwise one when *Θ*_sign_ = 1 (Figs 3–6C). In the next place, the *second* neighbor from the stimulus—next to the *first* in the same detour—took the smallest or second smallest |*B*_*α*_|. Then, the *third* neighbor of the stimulus—next to the *second*—took the largest or second largest |*B*_*α*_| which was opposite in sign to that of the *first*. One exception was the seven-armed case when *Θ*_sign_ = 0 (Fig. 6A); the *second* (arm 3) and the *third* (arm 4) respectively had the fourth smallest and the third largest, probably due to the outlying trial shown at the row 1 of column 4 in Fig. S6. Replacing the ordinary cases’ values with actual movements, the *first* actively pushed in the direction of the stimulated arm, while the *third* actively pushed oppositely to the *first*. These movements could make the *second* face forward, which indeed corresponded to the ranges of *Θ* in all the cases (Figs 3–6A,C; see also Videos S1, S2).

**Fig. 3.**
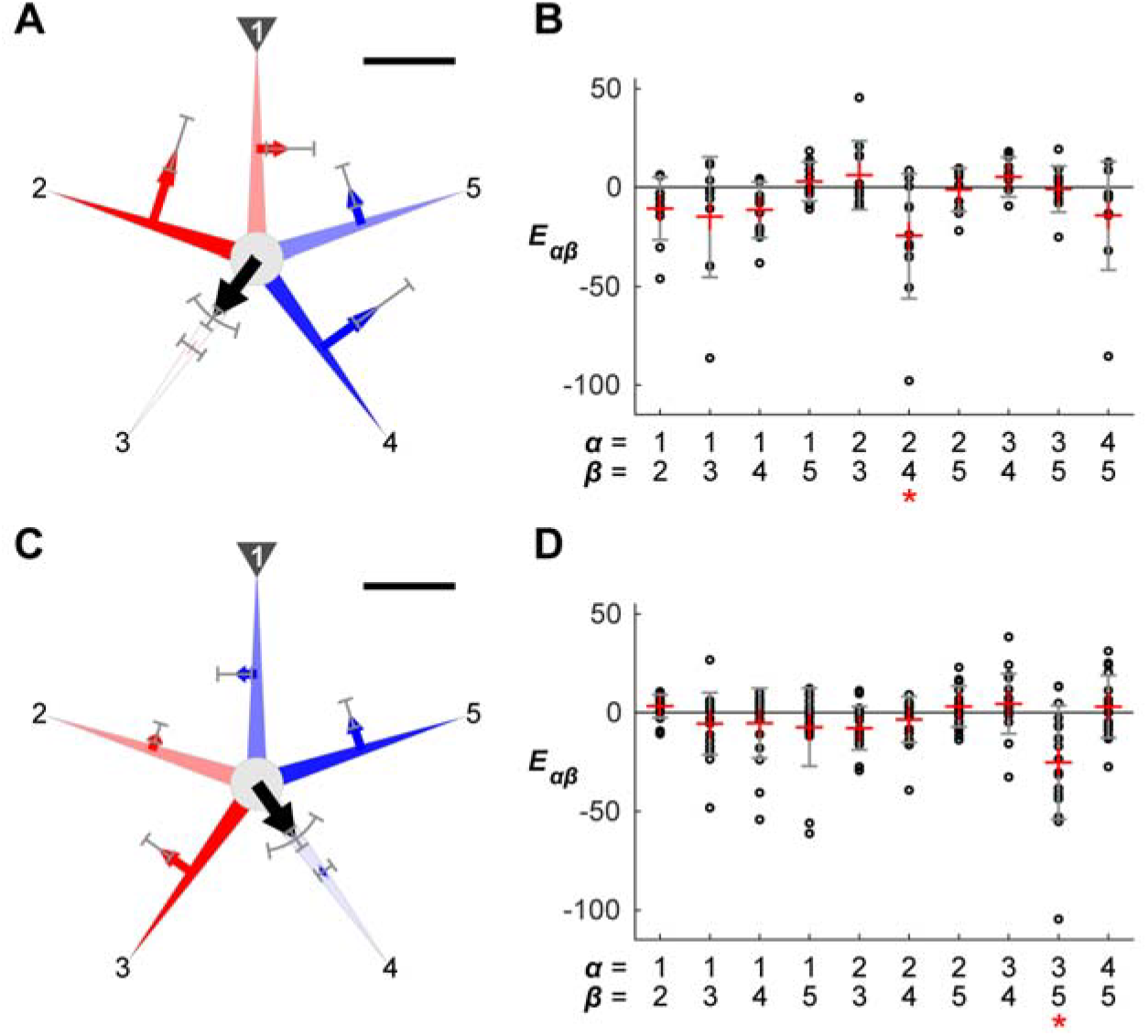
Five-armed locomotion grouped by moving direction in the brittle star *Ophiactis brachyaspis*. A,B: case where moving direction (*Θ*; c.f. Figs S1, S2) is angled clockwise from the opposite direction to the stimulated arm, i.e. *Θ* is negative and *Θ*_sign_ = 0 (eight individuals, 11 trials). C,D: case where *Θ* is positive (angled clockwise), i.e. *Θ*_sign_ = 1 (10 individuals, 19 trials); an exemplary locomotion in this case is shown in Video S1. A,C: schematized brittle stars reflecting the resultant quantitative values. Black arrows at the disks represent the measured means of moving distance (*S*; c.f. Fig. S1) by length and the measured means of *Θ* by angle. Error bars parallel to the disks’ arrows show *S*’s standard deviation (s.d.) and arc-shaped error bars represent *Θ*’s s.d. in data. The blue or red arrow at each arm represents the degree of being a left or right rower (*B*_*α*_; c.f. Figs S1, S2), reflecting the absolute median of each posterior mean by arrow length and the median of each posterior s.d. by error bars. When a posterior mean was negative/positive, its blue-leftward/red-rightward arrow extends from its arm, indicating that the arm pushed leftward/rightward (anticlockwise/clockwise), respectively. In each panel, the arm with the maximum absolute value in posterior mean is colored with the most vivid blue/red, while the other arms show lighter blue/red corresponding to the relative values to the maximum. Scale bars represent 40 mm for *S* and 50 for *B*_*α*_. B,D: degree of synchronization between two arms (*E*_*αβ*_ for the *α*- and *β*-th arms). Small circles represent measured values. Pair-by-pair red pluses indicate the medians of posterior means while error bars show the medians of posterior s.d. parameters. Negative/positive values represent that the paired movement of the *α*- and *β*-th arms synchronized in the opposite/same direction, respectively. Each asterisk indicates the pair with the negatively largest estimated mean, showing remarkable antiphase synchronization. All posterior distributions for both *B*_*α*_ and *E*_*αβ*_ were estimated under a better performed model in terms of WAIC, where *Θ*_sign_ is an explanatory variable for the mean and s.d.

**Fig. 4.**
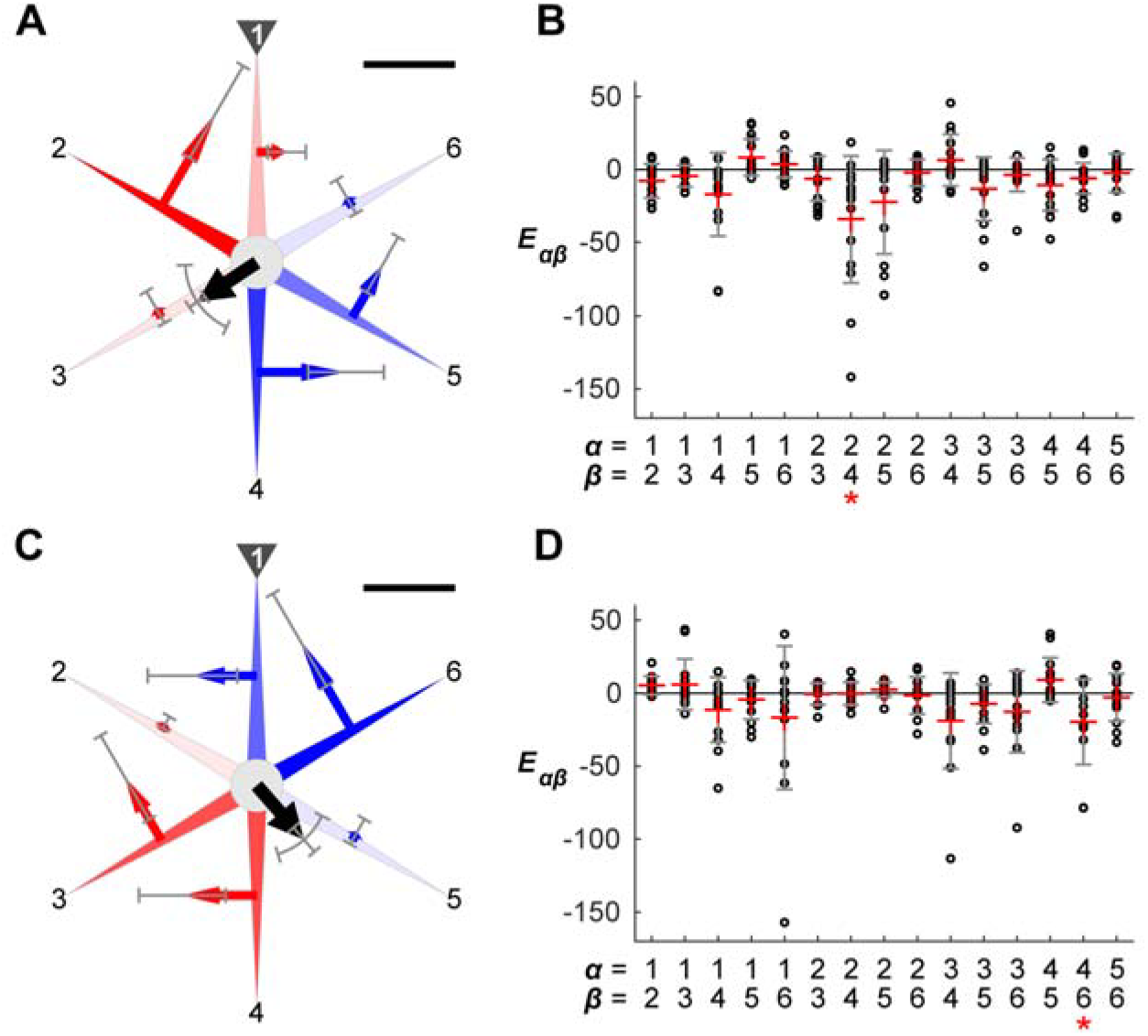
Six-armed locomotion grouped by moving direction in the brittle star *Ophiactis brachyaspis*. A,B: case where *Θ*_sign_ = 0 (eight individuals, 16 trials). C,D: case where *Θ*_sign_ = 1 (eight individuals, 14 trials); an exemplary locomotion in this case is shown in Video S2. A,C: schematized brittle stars reflecting the resultant quantitative values, as explained in Fig. 3. B,D: degree of synchronization between two arms (*E*_*αβ*_ for the *α*- and *β*-th arms), as explained in Fig. 3.

**Fig. 5.**
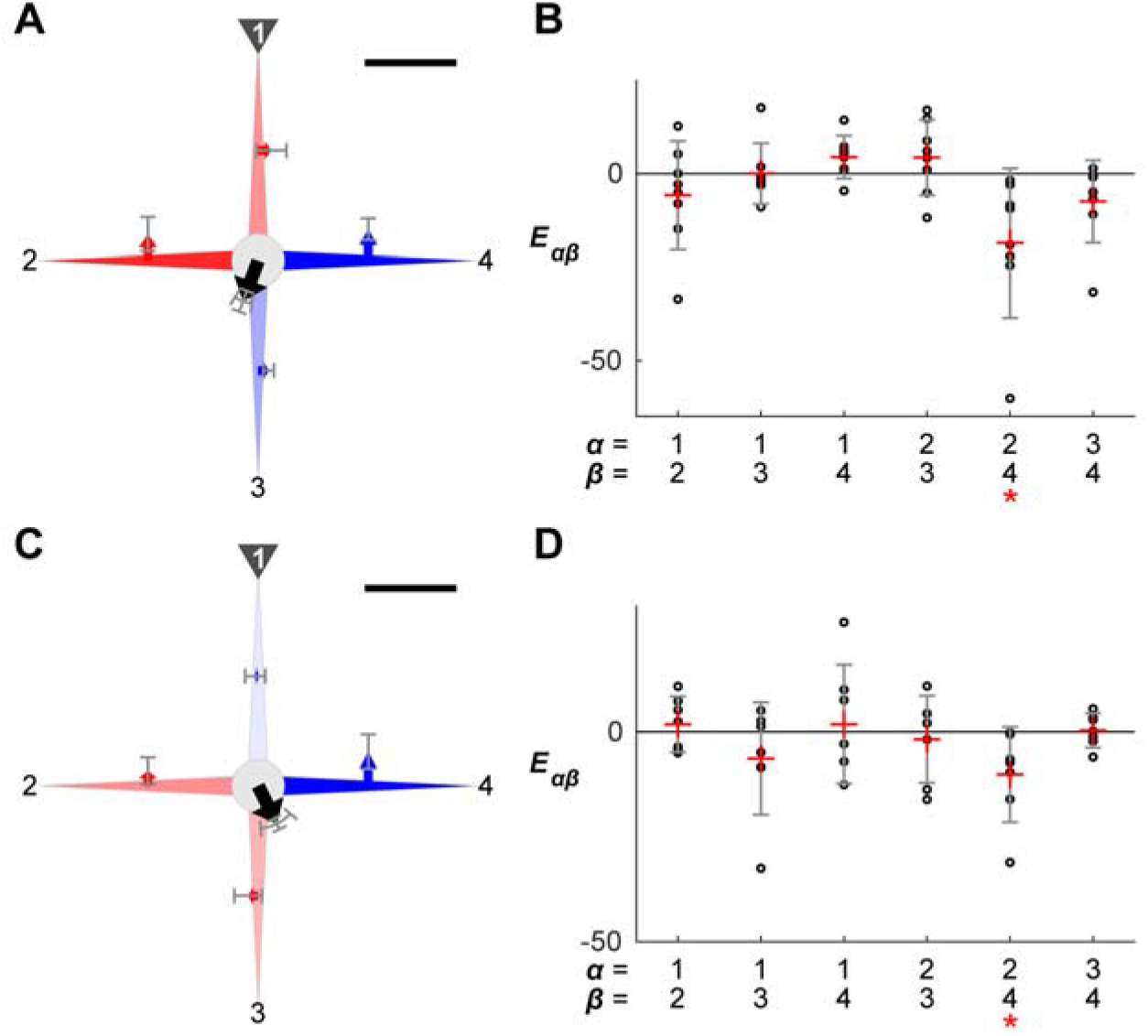
Four-armed locomotion grouped by moving direction in the brittle star *Ophiactis brachyaspis*. A,B: case where *Θ*_sign_ = 0 (one individuals, eight trials). C,D: case where *Θ*_sign_ = 1 (one individuals, seven trials). A,C: schematized brittle stars reflecting the resultant quantitative values, as explained in Fig. 3. B,D: degree of synchronization between two arms (*E*_*α*β_ for the *α*- and *β*-th arms), as explained in Fig. 3.

**Fig. 6.**
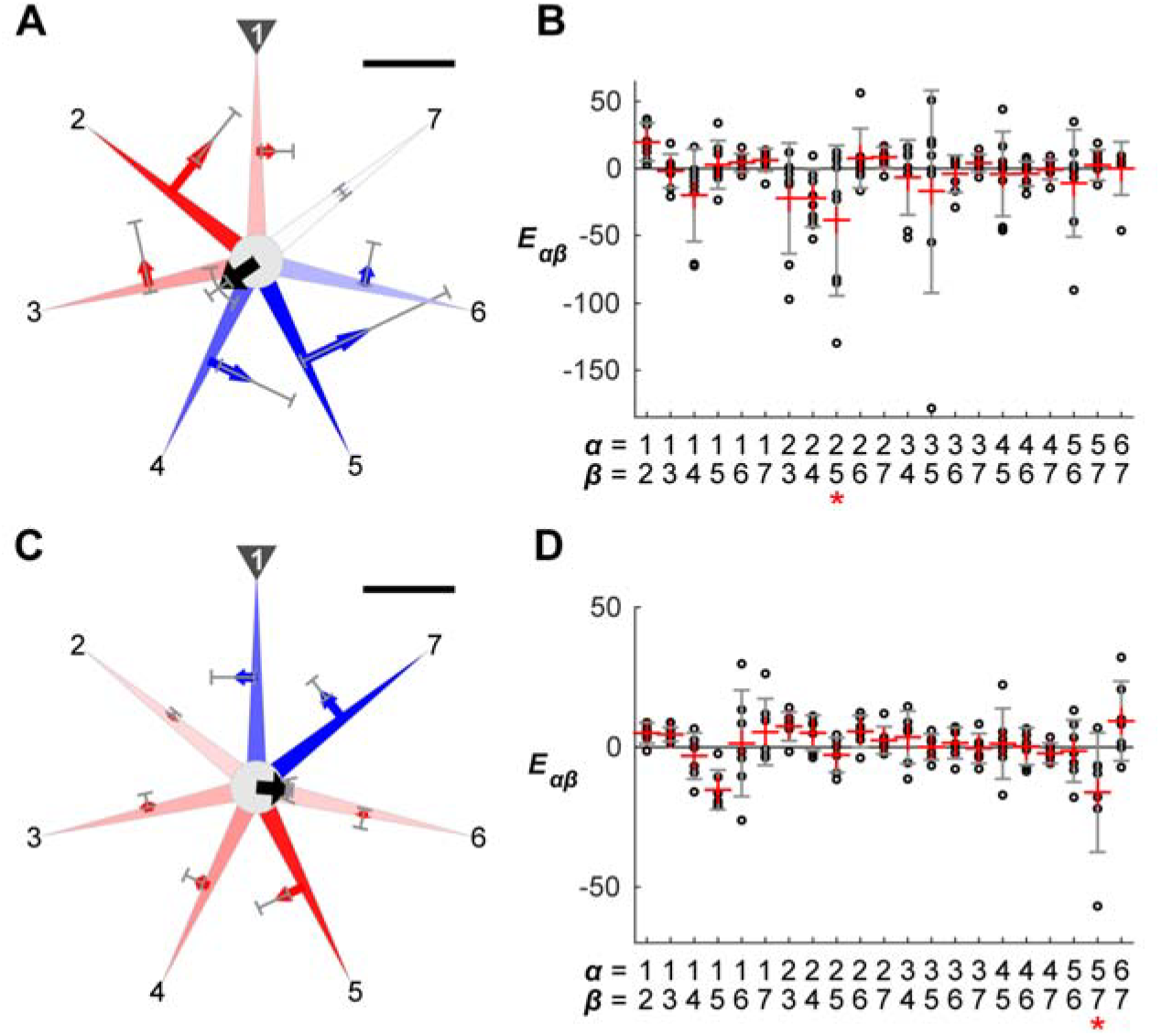
Seven-armed locomotion grouped by moving direction in the brittle star *Ophiactis brachyaspis*. A,B: case where *Θ*_sign_ = 0 (one individuals, eight trials). C,D: case where *Θ*_sign_ = 1 (one individuals, seven trials). A,C: schematized brittle stars reflecting the resultant quantitative values, as explained in Fig. 3. B,D: degree of synchronization between two arms (*E*_*αβ*_ for the *α*- and *β*-th arms), as explained in Fig. 3.

### Synchronization between two arms (*E*_*αβ*_)

The higher explanatory power of *Θ*_sign_ could also apply to the instance of the degree of synchronization between the *α*- and *β*-th arms, *E*_*αβ*_ (Eqn 4 in “Materials and methods”), because five-, six-, and seven-armed animals each brought the smallest WAIC in the model assuming *Θ*_sign_’s effect (Table 1). In the four-armed case, the model without an explanatory variable best performed while the presence of *Θ* or *Θ*_sign_ resulted in similar predictabilities. Accenting the significance of *Θ*_sign_ as with *B*_*α*_’s situation, we here show the resultant values of *E*_*αβ*_ discretely by the sign of *Θ*.

A side-by-side comparison with the *Θ*_sign_-based results of *B*_*α*_ shows us that the pair of the *first* and *third* rowers counting from the stimulus had the negatively largest medians of *E*_*αβ*_’s posterior means in most cases (Figs 3–6B,D). Although one exception was found in the seven-armed with *Θ*_sign_ = 0, the pair’s value *E*_24_ leaned negatively as well (Fig. 6B). These values gave a quantitative indication that these two arms tended to simultaneously push in the opposite direction, regardless of the number of arms.

## Discussion

Our study newly described the post-stimulus locomotion of brittle stars in a comparative context of four-, five-, six-, and seven-armed intact individuals in a single species. For this purpose, not stereotyping a discrete role of each arm, we introduced a quantitative index which can visualize each arm’s degree of being a left or right rower, namely *B*_*α*_. Coupled with other supportive values, we figured out a common rule accommodating different numbers of arms (Fig. 7). Our behavioral model would bring a general scheme of how a radially symmetrical animal maps ‘left and right’—in the same time, ‘front and back’—on its behavior via a decentralized control.

**Fig. 7.**
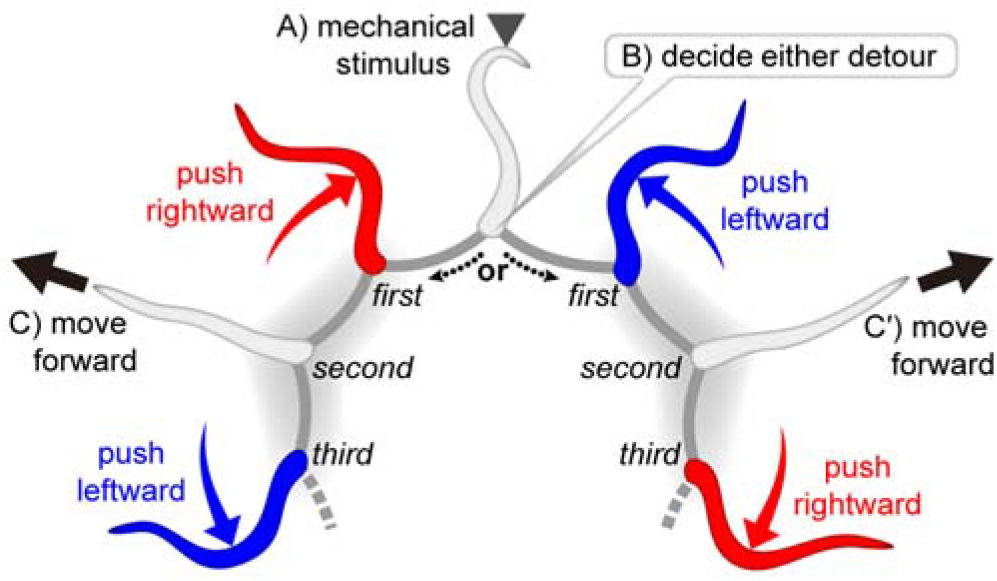
Model of how aversive tactile stimulation makes arm-by-arm locomotive movements in brittle stars with a flexible number of arms. The stimulated arm makes an afferent signal—(A)—which chiefly transfers in either of the clockwise or anticlockwise detour through inter-arm connections represented by the circumoral nerve ring. Which detour the signal dominates is determined by some perturbation—(B). Subsequently, one of the *first* neighboring arms to the stimulated arm pushes actively in the stimulus direction, while the *third* neighbor in the same detour pushes oppositely to the *first*. As a result, the *second* arm between the *first* and *third* faces forward in behavioral terms—(C) or (C′).

### Locomotor modes

In past quantitative studies using five-armed brittle stars, antiphase synchronization of two distant arms has been supported by assessing the stop and start timing of arm movements (Astley, 2012) and by evaluating *E*_*αβ*_ as in our study (Kano et al., 2017). This locomotor mode, which is referred to as “breast stroke” or “rowing,” is characterized by a leading arm and its side rowing arms (Romanes and Ewart, 1881; Preyer, 1887; Glaser, 1907; Arshavskii et al., 1976a,b; Astley, 2012; Kano et al., 2017; Clark et al., 2019). Our study figured out that the triplet of left-front-right could appear even in four-, six-, and seven-armed individuals, suggesting that this locomotor mode is determined anteriorly, not laterally or posteriorly. In addition, the two back arms in the five-armed locomotor mode have been often interpreted as passively dragged ones (Romanes and Ewart, 1881; Preyer, 1887; Arshavskii et al., 1976a,b; Watanabe et al., 2011); nevertheless, our study showed that these arms rather worked as weaker rowers since their values of *B*_*α*_ ranged either negatively or positively (Fig. 3A,C). In six- and seven-armed ophiuroids, back arms following the two strong rowers similarly exhibited a trend of rowers, whereas the backmost ones were usually neutral as to the leftward or rightward bias just like the leading arm (Figs 4A,C, 6A,C). Thus more arms could take charge of ‘rowers’ especially when a brittle star has more arms.

Although “breast stroke” or “rowing” is a frequently reported locomotor mode in five-armed brittle stars, some studies have also described patterns where there is no leading arm. One is called as “paddling” or “reverse rowing,” where a backmost arm is dragged while the other four actively row (Preyer, 1887; von Uexküll, 1905; Glaser, 1907; Arshavskii et al., 1976a; Astley, 2012). Such patterns without leading arms have been observed in free movement without experimental stimuli (Arshavskii et al., 1976a; Astley, 2012) as well as in escape behavior for a short time (Yee et al., 1987). In our study using *Ophiactis brachyaspis*, each trial seldom showed such a non-leading pattern (Figs S3–S6). Assuming this brittle star actually switches different locomotor modes, non-leading patterns might be employed only for several seconds after stimuli. In this case, our study might overlook or underestimate this urgent phase since we evenly analyzed one-minute duration after the beginning of the disk’s movement. In either case, as far as post-stimulus locomotion was quantified for the fixed period, it seems the locomotor mode with a leading arm is more usual in the intact individuals of the *Ophiactis* species regardless of how many arms they have.

### Decision of moving direction

Since brittle stars show no consistent front in behavioral terms as in most echinoderms, every arm can be a leading arm. Astley (2012) described their turning behavior in a short-term series, which was made by changing the roles of arms, not by rotating their body axis. As to an escape situation, several studies have observed that brittle stars avoid open or bright spaces (Cowles, 1910; Matsuzaka et al., 2017), a predator extract (Yee et al., 1987), and a KCl solution (Clark et al., 2019). However, few have probed into how such repellents make a certain reaction per arm to decide the moving direction of a whole individual. Since light and liquid diffuse in water, it is difficult to stimulate only a single target arm. Especially for small brittle stars such as *Ophiactis* species, tactile stimulation would perform better with the aim to understand how signals from a stimulated arm affect the movements of the other arms.

In our study, two quantitative indices calculated from the filtered angular velocity of arms—*B*_*α*_ and *E*_*αβ*_—and one obtained from the original coordinate data— *Θ*—could together visualize the ophiuroid locomotion without contradiction (Figs 3–6). Postulating each average of the two *Θ*_sign_-based patterns as a representative, our numerical results suggest the most frequent locomotion pattern after aversive tactile stimulation: a leading arm emerges at the *second* neighbor of a stimulated arm, while side arms adjacent to the leader synchronously push backward, no matter how many arms a brittle star has. To realize this bilateral distribution with a high probability, it can be assumed that an afferent signal from an arm makes one of the *first* neighboring arms be an active rower which pushes in the direction of the signaling arm, the *second* neighboring arm be an inactive one which has a less directional preference, and the *third* neighboring arm be another active one which pushes synchronously but oppositely to the *first*’s pushing (Fig. 7). Accordingly, the *second* faces forward while the *first*, *third*, and some rear arms work on its both sides. In this model, whether the clockwise or anticlockwise *second* arm becomes a leading arm depends on in which detour the signal from the stimulated arm dominantly transfers, which is determined by some perturbation. It seems the signal never makes symmetrical effects on both detours, contrary to our initial expectation that the stimulated arm’s both neighbors push toward the stimulus to let the disk escape just opposite to the stimulus.

Under our model shown in Fig. 7, brittle stars with more arms would have a more risk of ‘escape to a threat.’ If the front is placed ideally around the *second* neighboring arm from the stimulus, four-, five-, six-, and seven-armed animals will respectively show 0, 36, 60, and 77 deg in average |*Θ*|. In fact, the estimation from measured data copied it reasonably—17, 29, 46, and 70 deg, respectively—, and trials where moving direction rather inclined toward the stimulated arm (90 < |*Θ*| ≤ 180) were more frequent as a body had more arms: 0/15, 1/30, 3/30, and 5/15, respectively (Fig. 2). Although the ‘escape to a threat’ behavior is considered less adaptive, an evolutionary background would explain it. It has been proposed that primitive ophiuroids showed pentaradial symmetry (Paul and Smith, 1984; Sumrall and Wray, 2007), implying that brittle stars had developed a locomotion mechanism which worked optimally for the five-armed body. Some exceptional individuals in arm number, at least the four-, six-, and seven-armed bodies, probably have kept following this initial plan without vital issues. Meanwhile, escape direction could be more or less bent as a side effect, and the minority of four- and seven-armed ones might be a reflection of some inconvenience in control mechanism or its expression.

Our study has significance to understand how behavioral direction is expressed in a body without antero-posterior and left-right axes. Even when the individual body is round, some direction-making signal could transfer linearly in one direction at a local view (Fig. 7), just like a wave on a string or neural transmission in the spinal cord. Suppose brittle stars use this strategy, it seems not important how many segments with identical function are counted in the pathway. Otherwise, animal species would never allow individual difference in the number of motile organs.

### Inter-arm interaction

The inter-arm connection depicted in Fig. 7 is recognizable as the circumoral nerve ring—the main nervous system running in the disk. This correspondence is indicated by its orbital morphology as well as previous experimental support for the importance of the nerve ring in locomotion (Mangold, 1909; Diebschlag, 1938; Arshavskii et al., 1976a,b; Matsuzaka et al., 2017; Clark et al., 2019). Although it is difficult to measure neural activity in the small body of *Ophiactis* species, such an internal neural network can be suggested from behavioral modeling based on external observation. Given the simplicity of the ophiuroid nervous system, we can assume each arm’s movement directly reflects neural activity in each unit, which could be explained even by a couple of neurons. For instance, the observed locomotion would be a useful material for testing “neuron ring” models (Suzuki et al., 1971; Matsuoka, 1985) to know how circularly arranged neurons work in the real world. Taking advantage of the unique individual difference in fissiparous brittle stars, we are able to demonstrate them with different neuron numbers as in computer, which would build a new bridge between theoretical biology and experimental biology.

Besides the crucial role of neural interactions, Kano et al. (2017) found the ophiuroid’s ability to immediately change their locomotion patterns after the loss of certain numbers of arms, and then built an ophiuroid-like robot which imitated the adaptive locomotion via a local feedback without any preprogrammed control. Other robotics studies have also suggested the importance of ‘physical’ interactions in movement coordination which is independent to electrical circuits (Owaki et al., 2013; Owaki and Ishiguro, 2017). Taking account of these researches as well, it is not likely that four- to seven-armed individuals each employ a different central control system while counting the total number of arms. Each functional unit—e.g. each arm, each branch in the nerve ring—would just refer to the states of its nearest-neighbor units while ignoring distant ones; nevertheless, a coordinated pattern casually arises at a level of individuals, no matter how many units they own. In this perspective, the lower predictabilities of the individuality-assuming model (Table 1) imply that more important structural hierarchy for a brittle star might be each unit, rather than an individual body. A trial-by-trial variability in moving direction and other indices (Figs S3–S6) might reflect the influence of physical properties such as arms’ posture at each moment, although a circular neural network would chiefly design the average orientation, where the stimulated arm’s *second* neighbor faces forward (Fig. 7).

Although escape direction was unexpected in the more-armed cases, the resultant concept fits our initial hypothesis in terms of a decentralized design. High independence of each equivalent body sector must contribute to ophiuroid evolution that allows individual difference in appendage number. It may be a reason why some species such as *Ophiactis brachyaspis* have acquired fissiparity, being capable of drastic morphological changes in a life cycle while retaining its locomotive ability. For a potential application, unique ‘non-brained’ strategy of fissiparous brittle stars will be a hint of a highly flexible design in robotics.

## Acknowledgments

We thank Mr. Keita Harada in Shirahama Aquarium, Kyoto University, for sending us the fresh individuals of brittle stars from the aquarium. We are grateful to the member of Stan Community (https://mc-stan.org/community/) for suggesting better methods of statistical modeling.

## Data availability

The datasets generated and/or analyzed during the current study are available in the Figshare repository, https://doi.org/10.6084/m9.figshare.8019827.v3.

## Authors’ contributions

D.W. and H.A. designed the study and conducted behavioral experiments and measurements. D.W. and K.K. performed statistical modeling. All authors wrote the manuscript and approved the final manuscript.

## Supplementary material

**Video S1. Locomotion of a five-armed individual of the brittle star *Ophiactis brachyaspis*.** Quantitative analysis of this trial is presented in Fig. S2. Resultant values are schematized at the asterisked panel in Fig. S3.

**Video S2. Locomotion of a six-armed individual of the brittle star *Ophiactis brachyaspis*.** Resultant values are schematized at the asterisked panel in Fig. S4.

**Fig. S1.**
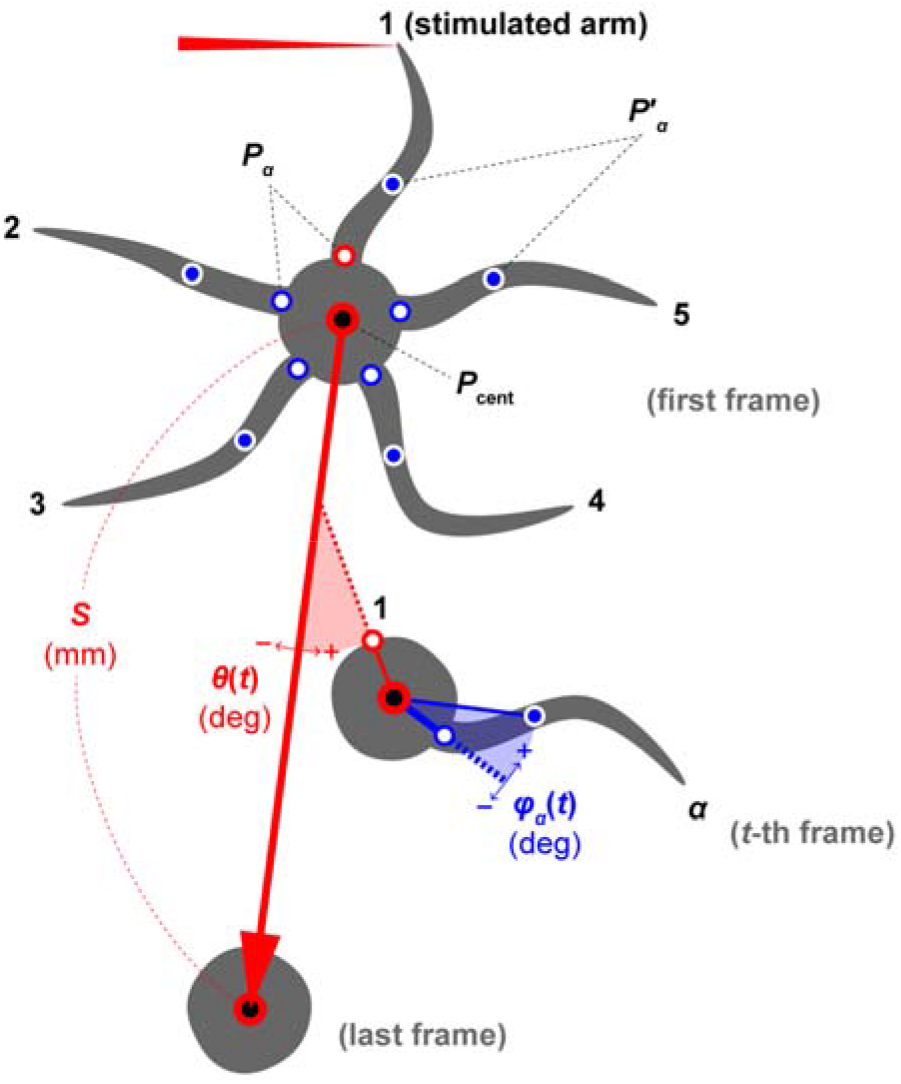
Measurements in the locomotion of the brittle star *Ophiactis brachyaspis*. Schematic five-armed brittle stars are shown at the first (*t* = 1), *t*-th, and last (*t* = 600) frames as an example. Not all arms are shown except for the first frame. The arm index, *α*, takes 1 to 5, where the stimulated arm is numbered 1. Blue-filled circles indicate the coordinate points of *P*′_*α*_(*t*) while open circles show those of *P*_*α*_(*t*). Particularly, *P*_1_(*t*) is indicated by red-lined open circles. The gravity of center of *P*_*α*_(*t*), namely *P*_cent_(*t*), is represented by red-lined filled circles. *φ*_*α*_(*t*) is the arm angle made by *P*_*α*_(*t*), *P*_cent_(*t*), and *P*′_*α*_(*t*). *θ*(*t*) is the angle made by the segment *P*_cent_(1)*P*_cent_(600) and the segment *P*_1_(*t*)*P*_cent_(*t*), representing the direction of the stimulated arm compared to moving direction. The moving distance *S* corresponds to the length of the segment *P*_cent_(1)*P*_cent_(600).

**Fig. S2.**
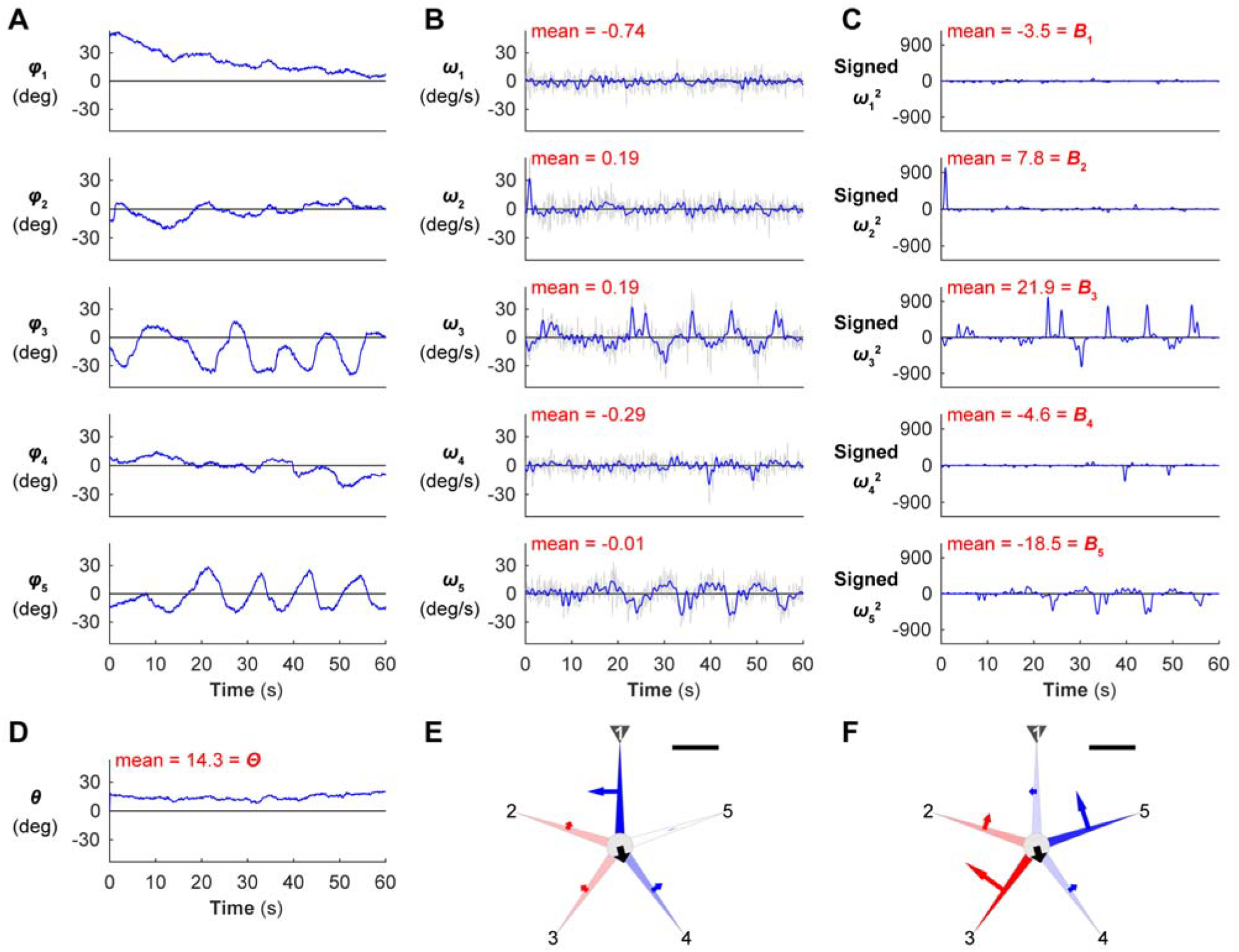
Calculation and visualization in a five-armed example of the locomotion of the brittle star *Ophiactis brachyaspis*. A: temporal change of φ_*α*_(*t*) deg (c.f. Fig. S1). B: temporal change of ω_*α*_(*t*) deg/s—angular velocity of φ_*α*_(*t*). Background gray plots represent the original data while thicker blue plots show low-pass filtered data. Each plot’s “mean” shows the mean value of the filtered ω_*α*_(*t*) for *t* = 1,‖, 600. C: temporal change of signed ω_*α*_(*t*)^2^. Each plots’ “mean” shows its mean value for *t* = 1,…, 600, corresponding to *B*_*α*_—the degree of being a left or right rower in the *α*-th arm. D: temporal change of *θ*(*t*) deg (c.f. Fig. S1). The “mean” shows its mean value for *t* = 1,…, 600, corresponding to *Θ* (deg)—moving direction. E: schematized brittle star reflecting the mean *ω*_*α*_(*t*) calculated in B and *Θ* in D. F: schematized brittle star reflecting *B*_*α*_ in C and *Θ* in D. In E and F, each gray arrowhead indicates the stimulated arm numbered 1, with the number followed anticlockwise in order. The angles of black arrows at the disks represent *Θ*. An arm with a negative/positive mean value extends a blue-leftward/red-rightward arrow, respectively, with its length corresponding to the absolute value of its mean. Compared to the mean values of the original ω_*α*_(*t*) in E, *B*_*α*_ in F well explains actual locomotion (c.f. Video S1). Note that *B*_*α*_ originally reflects a returning direction by its sign (positive *B*_*α*_ denotes anticlockwise returning), but its schematized arrow here indicates a ‘pushing direction’ for simply imagining force to the ground (positive *B*_*α*_ denotes clockwise pushing, so apparently opposing the sign in Fig. S1). Scale bars represent 1.0 for the mean ω_*α*_(*t*) in E and 20 for *B*_*α*_ in F.

**Fig. S3.**
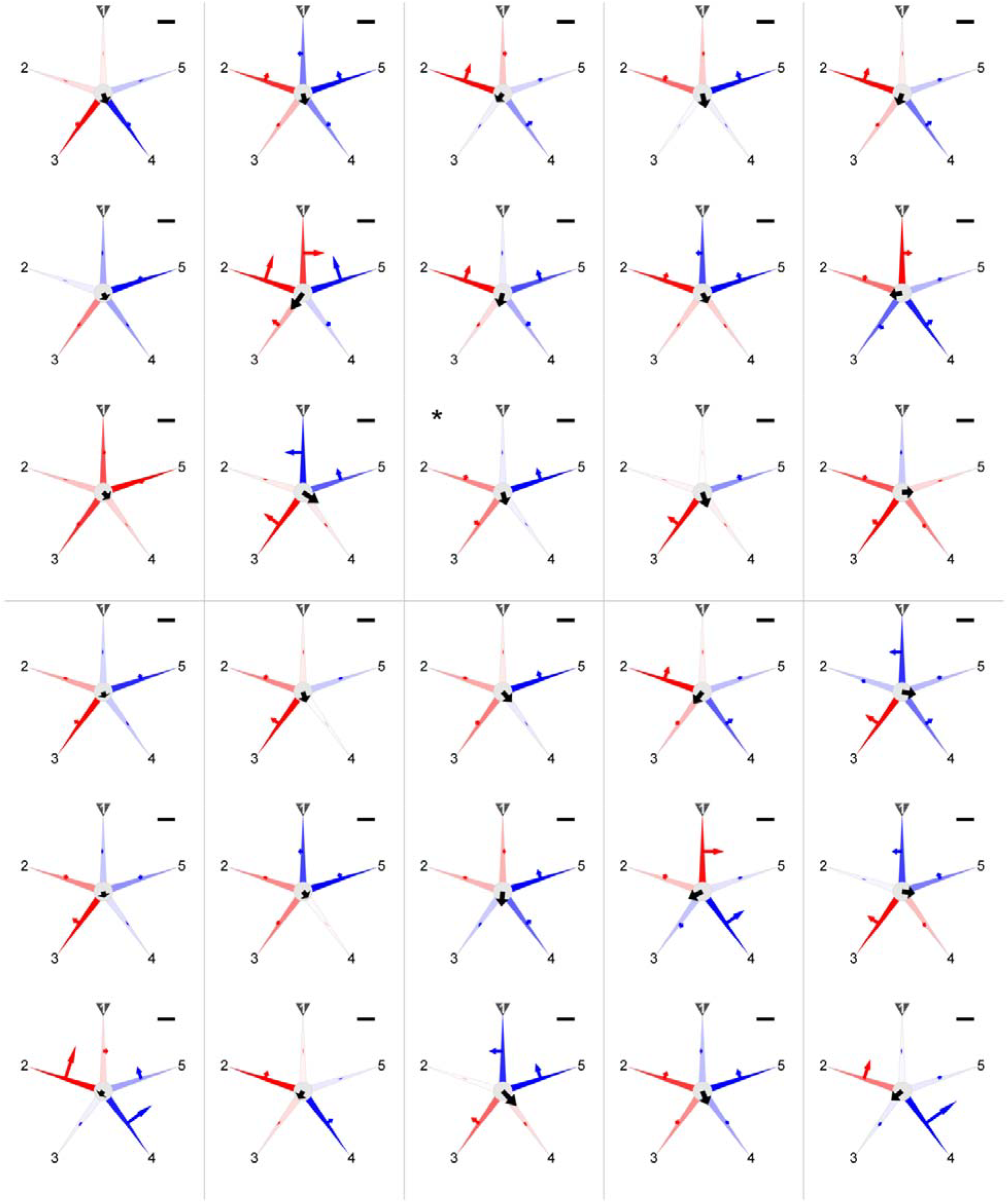
Five-armed trial-by-trial locomotion in the brittle star *Ophiactis brachyaspis*. Three trials were obtained from each of 10 individuals, which are partitioned by gray lines. Black arrows at the disks represent moving distance (*S*; c.f. Fig. S1) by length and moving direction (*Θ*; c.f. Figs S1, S2) by angle. An arm with a negative/positive value for the degree of being a left or right rower (*B*_*α*_; c.f. Figs S1, S2) extends a blue-leftward/red-rightward arrow, respectively, with its length corresponding to |*B*_*α*_|. In each panel, the arm with the maximum |*B*_*α*_| is colored with the most vivid blue/red, while the other arms show lighter blue/red corresponding to the relative values to the maximum. Scale bars represent 20 mm for *S* and 50 for *B*_*α*_. The asterisked trial (row 3, column 3) is shown in Video S1.

**Fig. S4.**
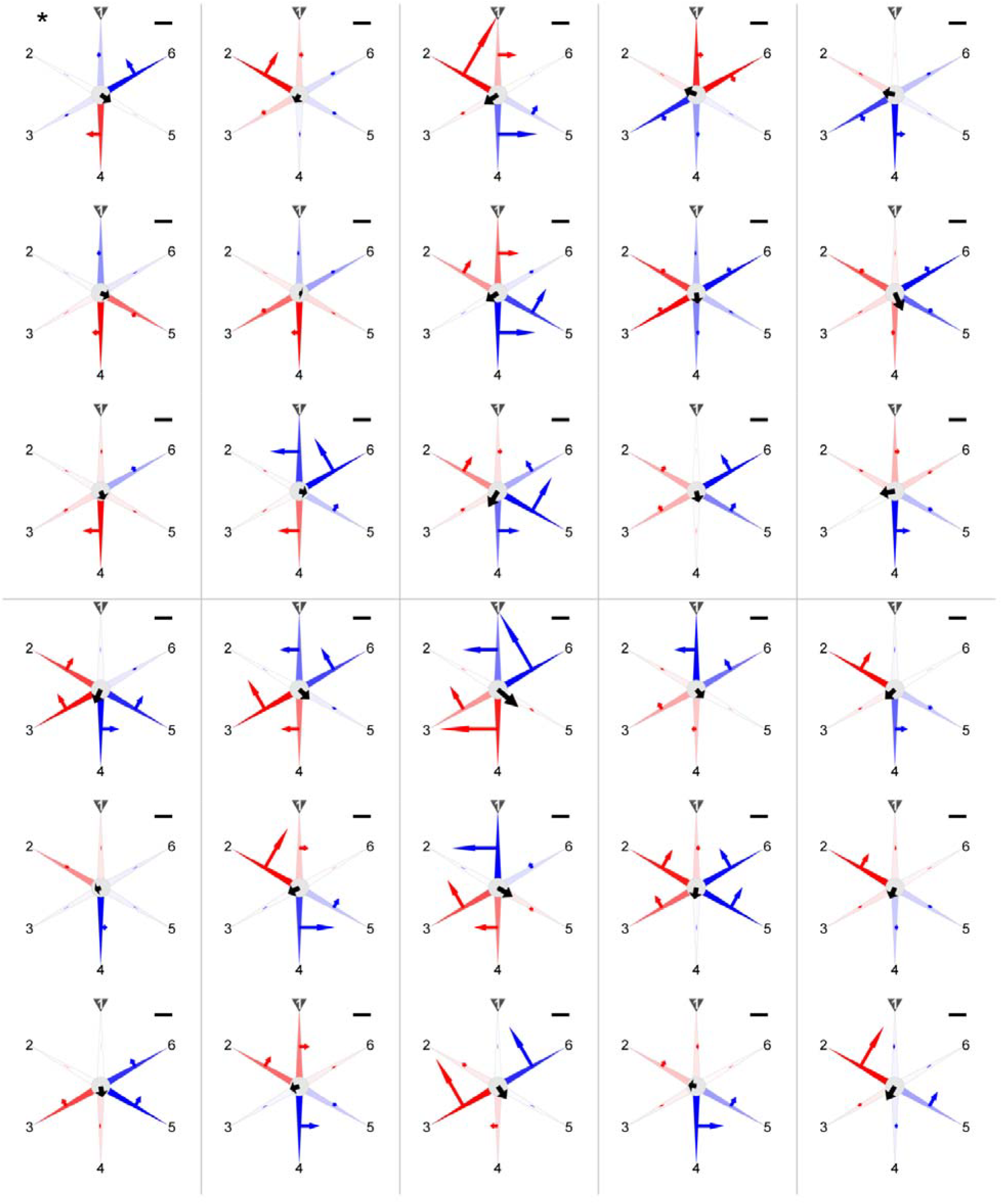
Six-armed trial-by-trial locomotion in the brittle star *Ophiactis brachyaspis*. Three trials were obtained from each of 10 individuals, which are partitioned by gray lines. Results are shown as in Fig. S3. The asterisked trial (row 1, column 1) is shown in Video S2.

**Fig. S5.**
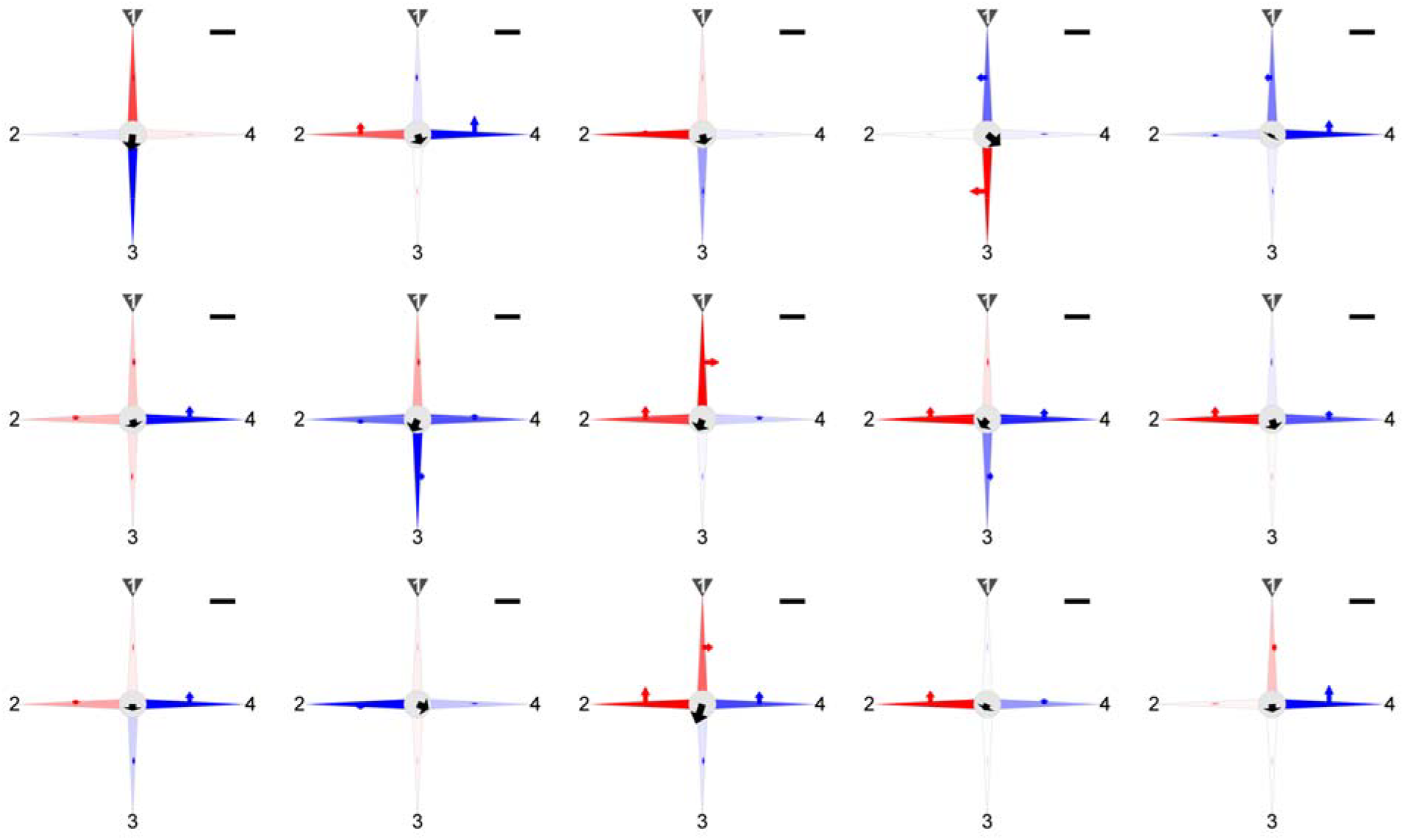
Four-armed trial-by-trial locomotion in the brittle star *Ophiactis brachyaspis*. Fifteen trials were obtained from one individual. Results are shown as in Fig. S3.

**Fig. S6.**
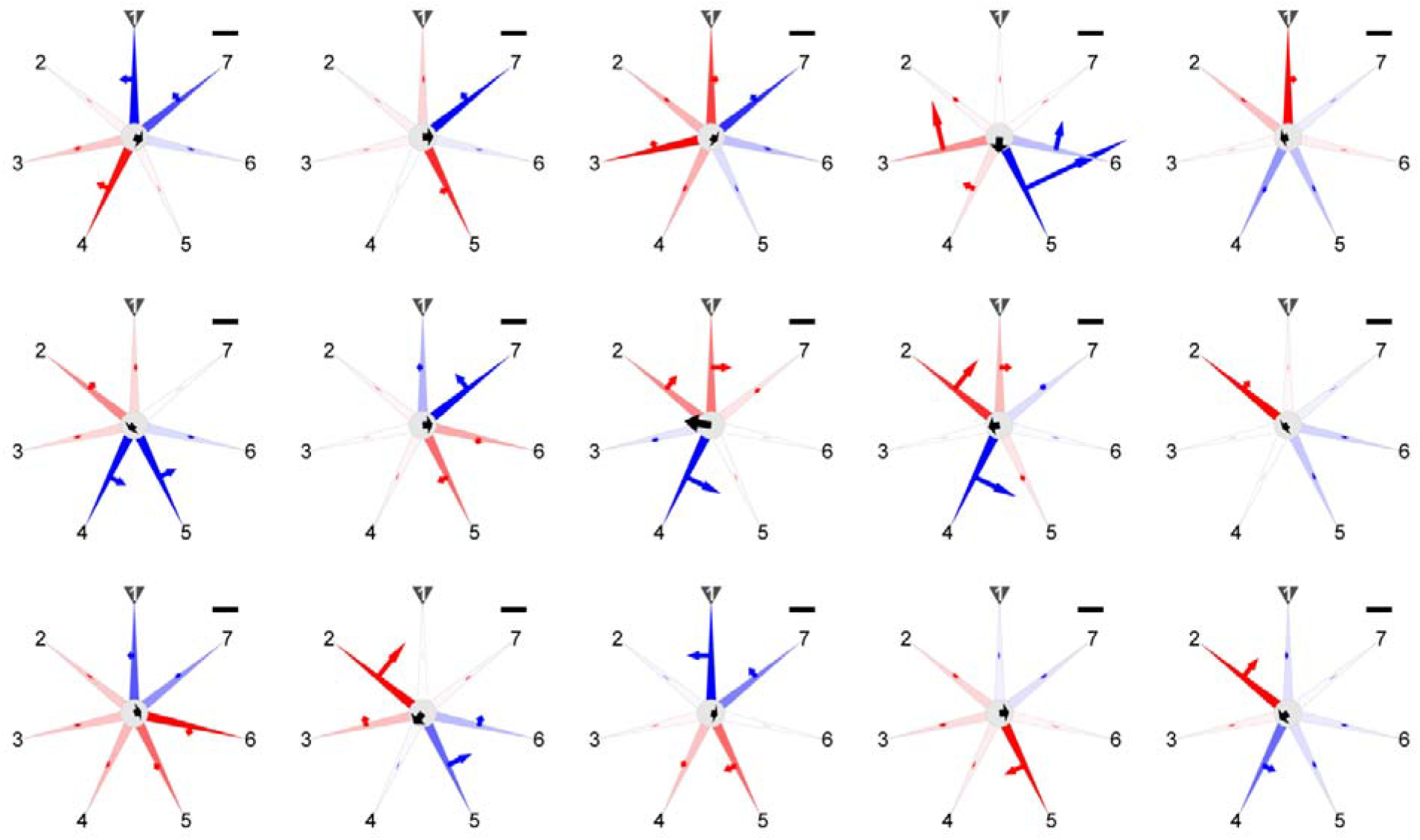
Seven-armed trial-by-trial locomotion in the brittle star *Ophiactis brachyaspis*. Fifteen trials were obtained from one individual. Results are shown as in Fig. S3.

